# A nuclear jamming transition in vertebrate organogenesis

**DOI:** 10.1101/2022.07.31.502244

**Authors:** Sangwoo Kim, Rana Amini, Otger Campàs

## Abstract

Jamming of cell collectives and associated rigidity transitions have been shown to play a key role in tissue dynamics, structure and morphogenesis. In cellular jamming, the physical state of the tissue is controlled by cellular density and the mechanics of cell-cell contacts. A potential contribution of subcellular organelles to the emergent tissue mechanics and architecture, as well as in the control of rigidity transitions, has not been explored. Here we show the existence of a nuclear jamming transition in which jamming of nuclei constrains cell movements beyond cellular jamming, with physical interactions between nuclei controlling the emergent physical properties and architecture of the tissue. We develop a computational framework and show that nuclear volume fraction and nuclear anisotropy are key parameters to understand the emergent tissue physical state. Analysis of tissue architecture during eye and brain development in zebrafish shows that these tissues undergo a nuclear jamming transition as they form, with increasing nuclear packing leading to more ordered cellular arrangements, reminiscent of the crystalline cellular packings in the functional adult eye. Our results reveal a novel rigidity transition associated with nuclear jamming, and highlight an important role for the nucleus in the control of emergent tissue mechanics and architecture.

---

Many developmental processes, including tissue morphogenesis and homeostasis, require a tight control of the tissue architecture and its physical state^1,2^. Ordered cellular packings play important functional roles in several organs, such as the inner ear^3^ or the eye^4,5^, whereas disordered cellular states are important for cell mixing at different developmental stages^6^. Similarly, the emergent mechanical state of the tissues (e.g., fluid/solid states) and its spatiotemporal control have been shown to play an important role in tissue morphogenesis^6–12^, cell dynamics^13–15^, cell differentiation^11,16^ and even tumor growth^17–19^. Connecting the emergent physical states at supracellular scales to the underlying processes that control them at cellular and subcellular scales is essential to understand how cells orchestrate embryonic development.

The physical state of embryonic tissues emerges from the collective physical interactions among cells. Mirroring inert systems, such as colloidal glasses, foams and emulsions^20–22^, glassy dynamics and characteristics of cellular jamming as well as rigidity transitions have been reported in cell monolayers *in vitro*^18,23,24^. More recently, rigidity transitions (jamming/glass transitions, fluid-to-solid transitions or tissue fluidization) have been observed *in vivo* within developing embryos and shown to play important functional roles^6,8,11,12,16,25,26^. Previous theoretical works have described the dynamics of cell collectives using various methodologies and predicted several observed behaviors, including different rigidity transitions^7,27–35^. To describe physical interactions among cells, these studies accounted for the structures that are generally thought to mediate mechanical interactions between cells, namely cortical tension, cell adhesion and traction forces, but neglected any organelles. While most organelles do not directly contribute to the mechanical interactions between cells, large ones could potentially play an important role in the emergent physical state of cell collectives and, specifically, of embryonic tissues.

The cell nucleus is the largest organelle in cells and is considerably stiffer than other cellular structures^36^. In tissues with cells much larger than their nuclei, the emergent mechanical state of the tissue is dominated by cell surface mechanics. However, during embryonic development, cells in different tissues and organs feature varying cell and nuclear sizes, with some cells being very large compared to their nuclei and others having nuclei nearly as large as the whole cell (Fig. 1). Variations in the ratio between nuclear to cell volume, namely the nuclear volume fraction, occur throughout embryonic development, and abnormal changes have been associated with malignant transformations in some tissues^37,38^. These observations indicate that nuclear mechanical interactions may not be negligible in many tissues, as previously suggested for syncytial embryos^39–41^ or in conditions leading to local nuclear crowding^42,43^. More generally, it is unclear how collective cell dynamics and the emergent tissue physical state and architecture may be affected by mechanical interactions among nuclei.

**Figure 1.**
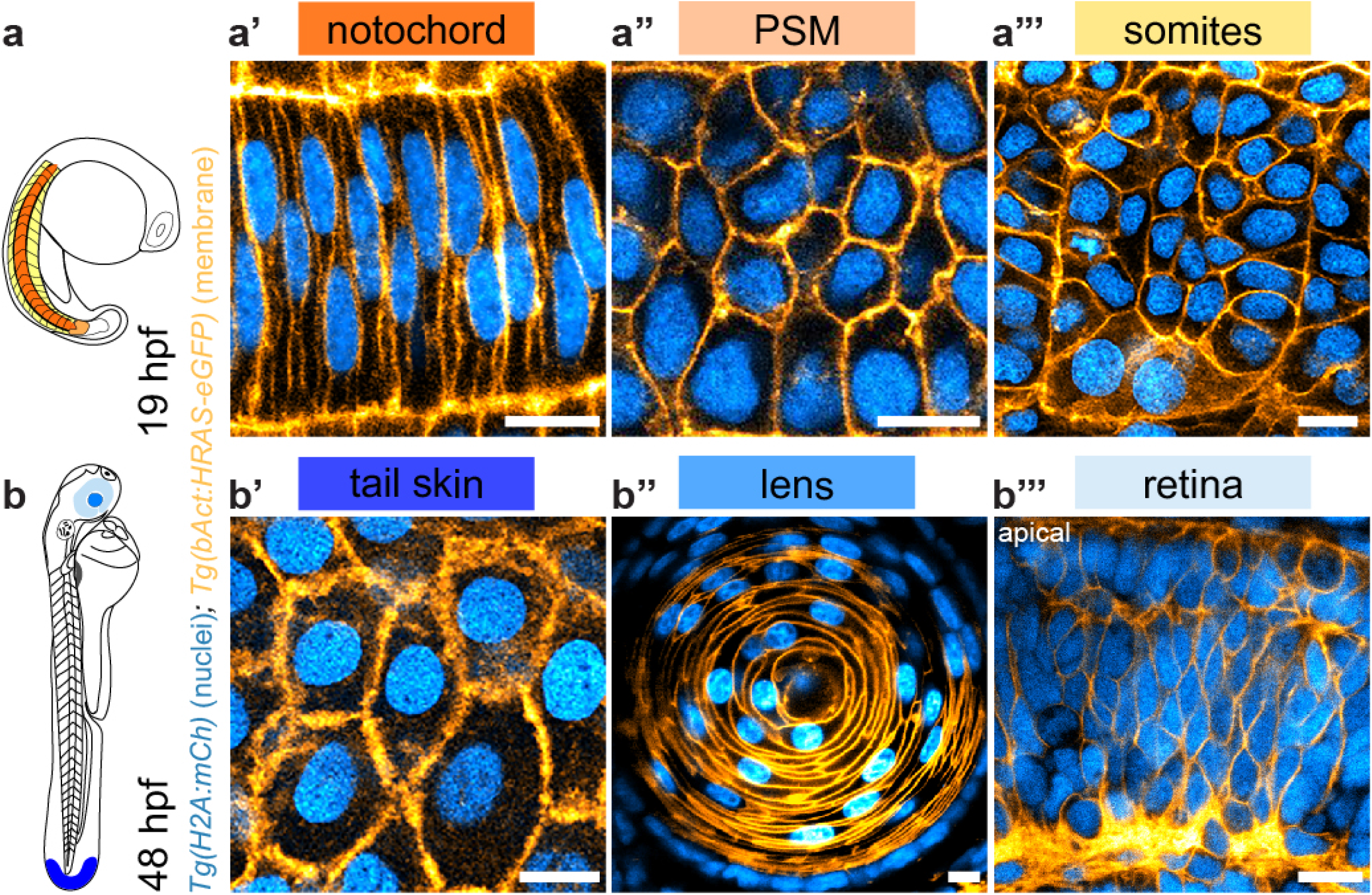
Varying nuclear volume fraction and anisotropy in various zebrafish tissues. **a-b**, Schematic representations of a zebrafish embryo at 19 (a) and 48 (b) hours post fertilization (hpf). **a’-a’’’** and **b’-b’’’**, representative confocal sections through different tissues of transgenic embryos with membrane (*Tg(bAct:HRAS-eGFP)*; orange) and nuclear (*Tg(H2A:mCh)*; cyan) markers at 19 hpf (a’, notochord; a’’, presomitic mesoderm (PSM); a’’’, somites) and 48 hpf (b’, tail skin; b’’, lens; b’’’, retina). Scale bars, 10 μm.

Here we study, both theoretically and experimentally, a new role for the nucleus in the control of emergent tissue mechanics and architecture. We develop a computational framework of tissue dynamics that accounts for both cell surface mechanics and the nucleus, and study the effect of varying nuclear volume fraction and anisotropy on the collective cellular dynamics and architecture of the tissue. Our results show that beyond a threshold value of the nuclear volume fraction, nuclei start affecting the tissue dynamics, mechanics and architecture, with the tissue becoming stiffer and displaying more ordered cellular packings. This nuclear jamming transition overtakes cellular jamming, restricting cellular movements and controlling many characteristics of the emergent tissue physical state. In vivo imaging of eye development in zebrafish shows that retinal tissues undergo a nuclear jamming transition as development progresses, causing a progressive increase in the order of cellular packings. We observe a similar transition during brain development, indicating that nuclear jamming is not unique to eye development and may constitute a more general mechanism to constrain cellular movements and establish ordered cellular packings in developing organs.

## Theoretical description

To understand the role of varying nuclear volume fraction and nuclear anisotropy in the emergent physical state of the tissue, we first developed a computational framework that accounts for both cellular and nuclear mechanical interactions (Fig. 2a-c). Building on the Active Foam description of tissue dynamics^7^, we introduced a nucleus within each cell as an individual, soft elliptical particle of fixed size *A_N_* and long and short semiaxis b and 𝑎, respectively, that determine the nuclear volume fraction *𝜙*_*N*_ ≡ *A_N_*/*A_C_* (with *A_C_* being the cell size) and nuclear anisotropy *𝛼_N_* ≡ *b*/*a* (Fig. 2a,b; Methods). Nuclei do not physically interact directly, but rather interact with the cell boundaries upon contact, with repulsive interactions between nuclei and cell boundaries defined by a repulsive harmonic potential (Fig. 2c; Methods). To reproduce the experimentally observed nuclear positioning close to the cell center^44,45^, we introduced a restoring force for the nucleus to the geometric cell center (Fig. 2c). While this effective restoring force enables realistic nuclei dynamics in cells, it does not qualitatively affect our results below.

**Figure 2.**
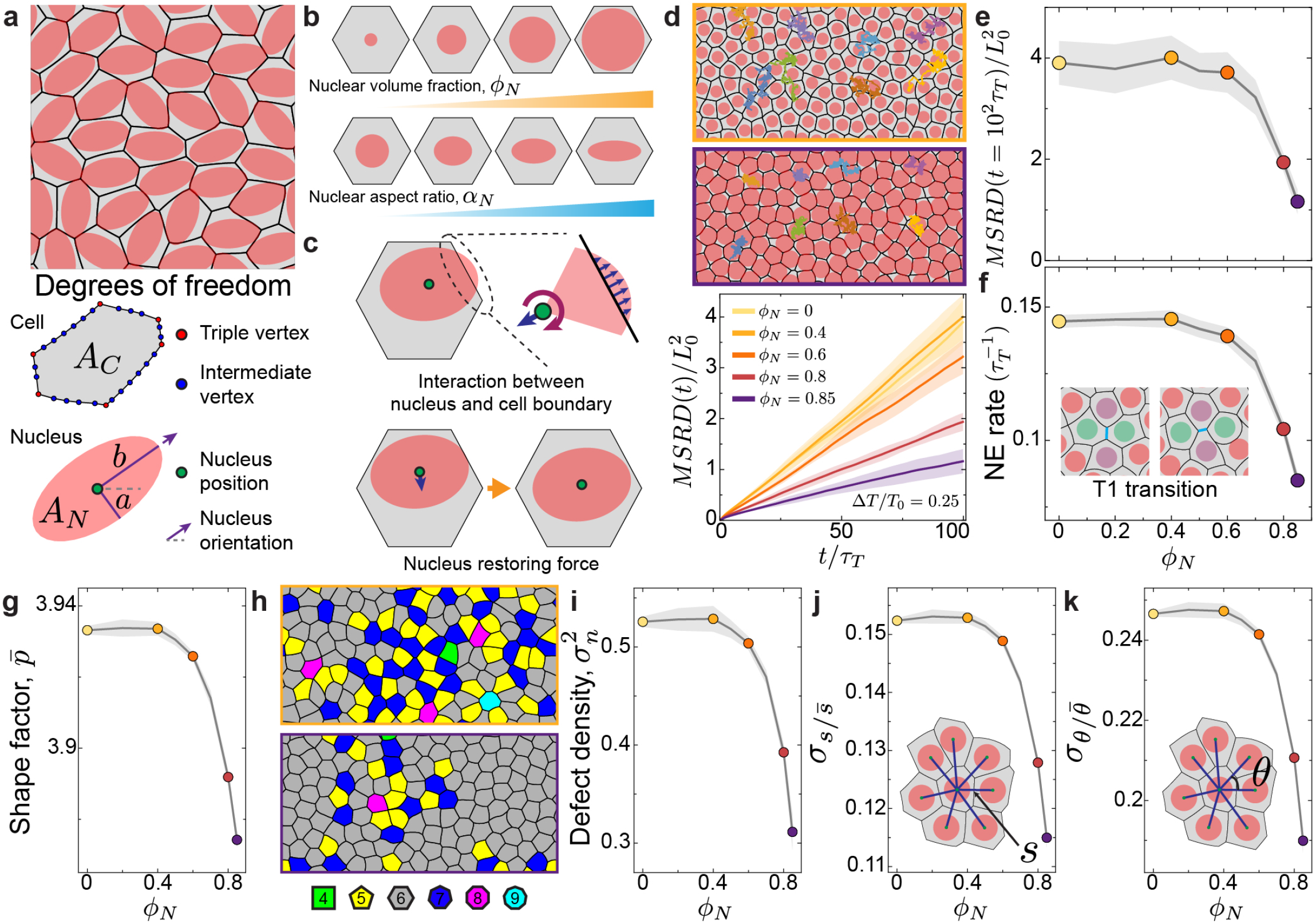
Characteristics of nuclear jamming for round nuclei. **a**, Snapshot of a simulated system configuration showing cells (black borders; gray interior) and nuclei (red), as well as the degrees of freedom for each cell and parameters characterizing nuclear size and shape. **b**, Schematic diagrams with examples of varying nuclear volume fraction *𝜙_N_* and nuclear aspect ratio *𝛼*_*N*_. **c**, Schematics showing the forces acting on the nuclei (Methods; Supplementary Information): force between the nucleus and the cell boundary (top), and effective nucleus restoring force (down). **d**, Representative cell trajectories for small (top, *𝜙_N_* = 0.4) and large (down, *𝜙*_*N*_ = 0.85) volume fractions. MSRD shows a progressive reduction in cell movements for increasing values of the nuclear volume fraction *𝜙_N_*. **e-k**, Long time MSRD values (e), neighbor exchange (NE) rate (f), shape factor 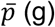, defect density 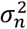 (i; snapshot of two systems configurations are shown in (h), color-coded in terms of number of neighbors), standard deviation of normalized bond length 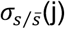, and standard deviation of normalized bond angle 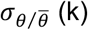 for different nuclear volume fractions. Error bands indicate standard deviation.

We first focus on the case of round nuclei *(𝛼_N_*= 1). For small nuclear volume fraction, cellular dynamics and tissue architecture are not affected by the presence of nuclei because these rarely interact (Fig. 2d). As the nuclear volume fraction surpasses a threshold 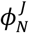 of approximately 0.6, all dynamic and structural measures show that cells start being more constrained (Fig. 2d-f; Supplementary Movie 1) and ordered (Fig. 2g-k) as a consequence of nuclear interactions. Both the normalized mean squared relative displacement 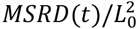, where *L*_*0*_ is the characteristic length scale of cell size; Methods; Fig. 2d,e) and the rate of cellular rearrangements (or T1 transition rate; Fig. 2f) decrease strongly as the nuclear volume fraction increases beyond the threshold 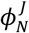 (Fig. 2d-f), indicating a rigidification of the tissue associated with nuclear jamming. The specific value of this nuclear jamming volume fraction is independent of the magnitude of active tension fluctuations at cell-cell contacts within the studied range (Extended Fig. 1). Very large values of the nuclear volume fraction nearly arrest the system even in the presence of substantial tension fluctuations, indicating that strong nuclear jamming can rigidify a tissue that would otherwise be fluidized by tension fluctuations (Fig. 2e). This is because nuclear jamming obstructs nuclear uncaging behavior even when cell-cell contacts undergo T1 transitions (Fig. 2f). For values of the nuclear volume fraction leading to nuclear jamming, the emergent tissue mechanics is controlled by the nuclear mechanics rather than by the mechanics of cell-cell contacts if nuclear stiffness *E_N_* is larger than the characteristic stress scale at cell-cell contacts, namely *T*_0_/*L*_0_ (with *T*_0_ being the average cell-cell contact tension; Methods). Indeed, cell movements and tissue dynamics are unaffected by the presence of nuclei if these are very soft (*E_N_* < *T*_0_/*L*_0_), but are strongly affected by stiff nuclei (*E_N_ ≫ T*_0_/*L*_0_) at large nuclear volume fractions (Extended Fig. 2).

Increasing the nuclear volume fraction above the nuclear jamming value 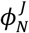 also affects the tissue architecture. For high values of the nuclear volume fraction, nuclei’s jammed states act as a geometric template for cell shape, so that cell shapes become more isotropic and their spatial arrangements more ordered. The shape factor, 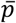, (namely the ratio of cell perimeter to square root of cell area; Methods; Fig. 2g), the topological defect density, 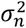 (as quantified by the variance of the topology distribution; Methods; Fig. 2h,i), as well as the variation in normalized bond lengths *s* and angles *𝜃* (quantified by the standard deviation of normalized bond lengths and angles, 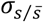 and 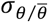 respectively; Methods; Fig. 2i,k), all decrease as the nuclear volume fraction increases,m indicating the formation of more ordered, crystalline cellular packing structure. Overall, these results indicate that high nuclear volume fractions for round nuclei lead to nuclear jamming, which overtakes cellular jamming, causing a strong reduction in cell movements, tissue rigidification and ordering of cellular architecture in the tissue.

Nuclear anisotropy can significantly change the resulting tissue dynamics and structure. For small nuclear anisotropy, namely *𝛼*_*N*_ ≲ 1.5, cell movements and rearrangement rates show the same qualitative behaviors as for round nuclei (Fig. 3a,b). However, for increasing values of nuclear anisotropy, the onset of nuclear jamming occurs at smaller values of nuclear volume fraction than for round nuclei, as shown by the reduction in both cell movements and neighbor exchange rates (Fig. 3a,b; Supplementary Movie 2). This is due to the fact that the long axis of nuclei start to interact with cell boundaries, as quantified by the mean number 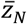 of contacts between nuclei and cell boundaries (Methods; Fig. 3c,d), enabling mechanical interactions at lower nuclear volume fractions. For values of the nuclear volume fraction and anisotropy leading to a long nuclear axis similar to the cell size, nuclei orient along opposed vertices in cells, as this is their optimal configuration within the constraints of cell shape (Fig. 3d). While this specific interaction between nuclei and cell boundaries does not strongly affect cell shapes (Fig. 3f), it hinders cell movements and rearrangement rates, indicating tissue rigidification (Fig. 3a,b). For even larger values of nuclear anisotropy and volume fraction, the highly anisotropic nuclei deform cell shapes, which become anisotropic (Fig. 3e,f,g). In these conditions, neighboring cells start to locally align and the system starts displaying local nematic order (Fig. 3i; Methods). This symmetry breaking leads to an enhancement in cell rearrangement rates that causes increased cell movements in the direction of the local nematic order (Fig. 3a,b). These results indicate that the interplay between nuclear jamming and nuclear anisotropy in the presence of active tension fluctuations can cause directional, collective cell movements.

**Figure 3.**
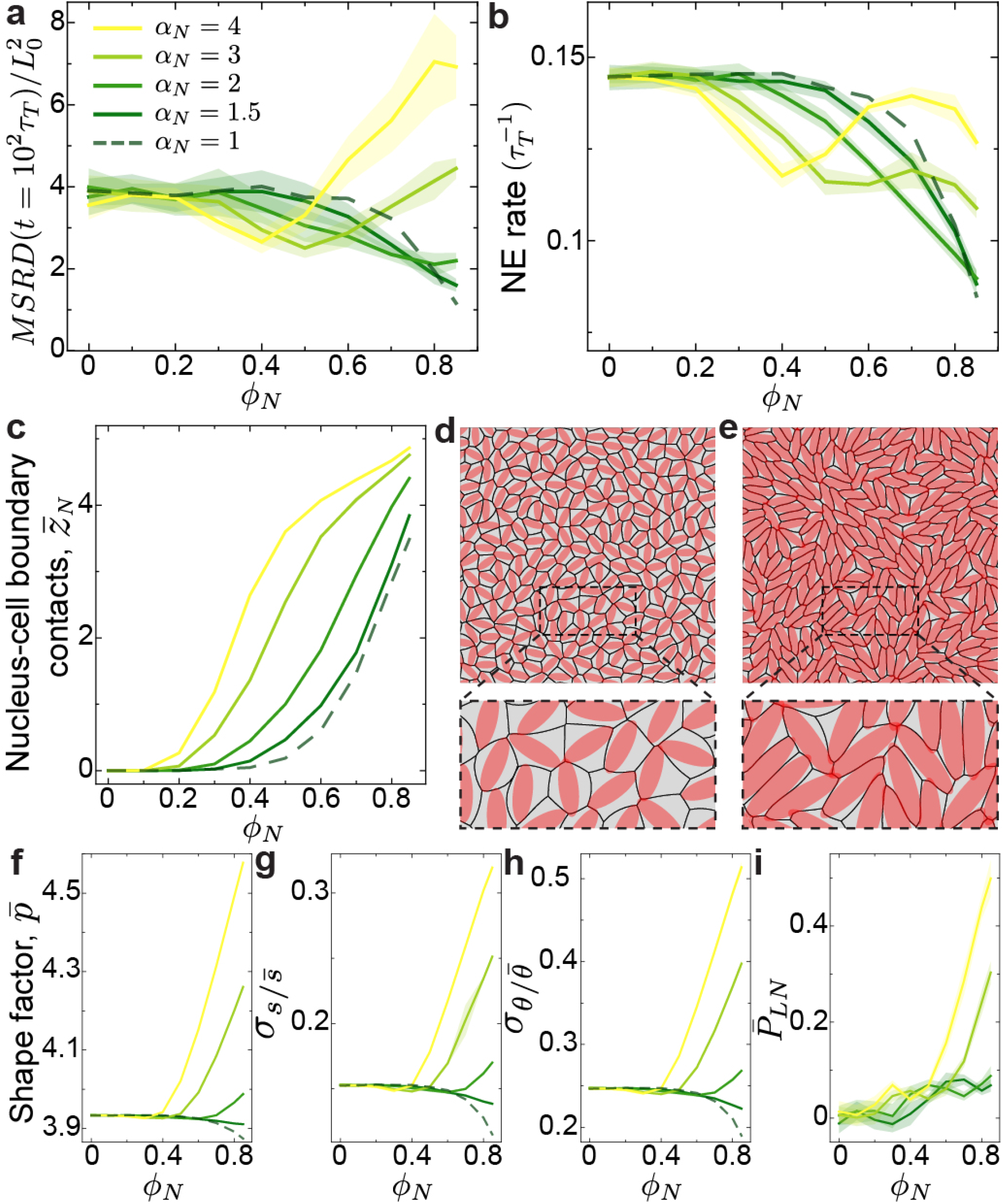
Role of nuclear anisotropy in nuclear jamming. **a-c**, Long timescale (*t*/*τ_r_* = 100) MSRD values (a), neighbor exchange rate (b), and mean number of contacts between nuclei and cell junctions 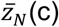 for different nuclear volume fraction (Methods). **d-e**, Simulated configurationsof anisotropic nuclear jamming (d; *𝜙_N_* = 0.5, *𝛼_N_* = 3) and local nematic ordering (e; *𝜙_N_* = 0.85, *𝛼_N_* = 4). Zoomed in configurations are shown for both cases (bottom). **f-i**, Shape factor 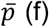 (f), standard deviation of normalized bond lengths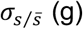 (g), standard deviation of normalized bond angle 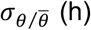, and local nematic order parameter 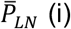 for different nuclear volume fractions (Methods). Error bands indicate standard deviation.

## Nuclear jamming during retina development

To determine if nuclear jamming plays an important role in living tissues, we imaged nuclei and cell boundaries (membranes) in different tissues of various organs at multiple stages of zebrafish development (Fig. 1a-b’’’). Among the imaged tissues, the retina displayed comparatively large nuclei compared to cell size (Fig. 1b’’’), which suggested the possibility of nuclear jamming occurring in this tissue. In vertebrates, out-pocketing of the neural tube gives rise to a single-layered neuroepithelium that defines the initial stages of the retina, with elongated progenitor cells attached to both apical surface and basal lamina^46^ (Fig. 4d, 24-36 hpf; Extended Figs. 3a and 4a,b). As development progresses, retinal progenitor cells detach from tissue boundaries at approximately 42 hpf and differentiate into retinal neurons^47^ (Extended Fig. 3b), creating a tissue with more compact cells (Fig. 4d, 55 hpf; Extended Fig. 4c). The retina is gradually transformed into a layered tissue in which cell bodies of its five different neuron types are positioned at three morphologically distinct layers from apical-to-basal: the outer nuclear layer (ONL), inner nuclear layer (INL), and the retinal ganglion cell layer (RGL)^48–50^ (Fig. 4d, 55-120 hpf; Extended Fig. 4c-f).

**Figure 4.**
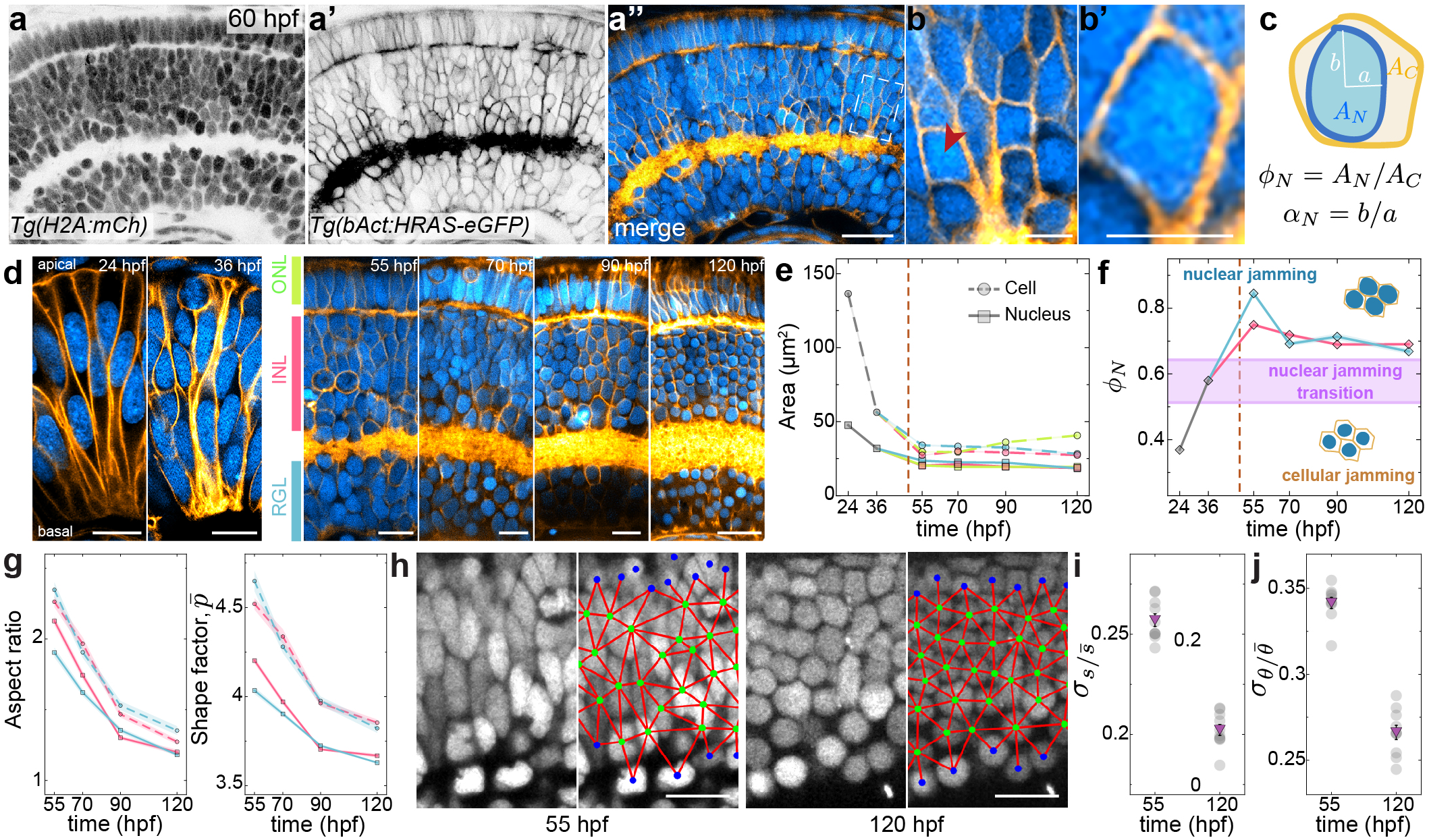
Nuclear jamming transition during zebrafish retina development. **a-a’’**, Representative confocal sections of a 60 hpf zebrafish retina (a, nuclear label, *Tg(H2A:mCh)*, cyan; b, membrane label, *Tg(bAct:HRAS-eGFP)*, orange; c, merge). Scale bar, 20 μm. **b**, Higher magnification inset of the outlined region in **a’’. b’**, High magnification of a cell (red arrowhead) in **b**. Scale bars in b-b’, 5 μm. **c**, Schematics of a retinal cell with the relevant definitions to quantify the nuclear volume fraction *𝜙*_*N*_ and the nuclear anisotropy *𝛼*_*N*_. **d**, Representative confocal sections of zebrafish retina at 24, 36, 55, 70, 90 and 120 hpf, with definitions of the distinct retinal layers (from apical to basal; ONL, INL, RGL) that gradually emerge after cell detachment from tissue boundaries (Extended Fig. 3). Scale bars, 10 μm. **e**, Temporal change in the area (*A_C_*; dashed lines) and nuclei (*A_N_*; solid lines) of retinal cells before cell detachment from tissue boundaries (24 and 36 hpf; vertical dashed line indicates approximate time of cell detachment from tissue boundaries) and in the different retinal layers (ONL, green; INL, pink; RGL; light blue) after cell detachment from the boundaries (55, 70, 90, 120 hpf). Statistics: 24 hpf (N=12, n=219), 36 hpf (N=7, n=319), 55 hpf (ONL: N=9, n=181; INL: N=14, n=470; RGL: N=8, n=195), 70 hpf (ONL: N=9, n=231; INL: N=9, n=342; RGL: N=8, n=275), 90 hpf (ONL: N=5, n=240; INL: N=6, n=386; RGL: N=5, n=150), 120 hpf (ONL: N=6, n=194; INL: N=7, n=294; RGL: N=6, n=151). N and n indicate number of samples and total analyzed cell/nuclei, respectively. **f**, Temporal evolution of nuclear volume fraction in the retina (only INL and RGL; Methods). Dashed vertical line indicates the approximate point at which cells detach from tissue boundaries. Horizontal violet band shows the nuclear jamming volume fraction predicted theoretically (Fig. 2). **g**, Temporal evolution of aspect ratio (left) and shape factor (right) for cells (dashed lines) and nuclei (solid lines) in the INL and RGL. **h**, Confocal section of the INL showing the nuclei (left) and the network (red) connecting the geometrical centers (green; blue indicate those at the INL boundary) of nearest neighbors (right; overlaid) at both 55 and 120 hpf. **i-j**, Variation in distances *s* (i) and angles *𝜃* (j) of nearest neighbors (Methods).

To investigate if the retina undergoes nuclear jamming, we quantified both the nuclear volume fraction *𝜙*_*N*_ and nuclear anisotropy *𝛼*_*N*_ as it developed (Fig. 4c). Analysis of cell and nuclear sizes revealed their substantial decrease over time (Fig. 4e, dashed line; Methods). While both cell and nuclear sizes decrease, cell size does so to a larger extent, causing a sharp increase in the nuclear volume fraction as cells detach from tissue surfaces. The nuclear volume fraction in both INL and RGL eventually crosses the threshold for nuclear jamming (Fig. 4f), at approximately 55 hpf, with both layers showing nuclei nearly as large as the cells themselves. While the ONL also shows a similar increase, we excluded it from the analysis because of the strong potential influence from the tissue boundaries in the nuclear jamming process. As development further proceeds and the retina progressively reaches its mature organization (70-120 hpf), the nuclear volume fraction in both INL and RGL decreases slightly but always remains above the nuclear jamming threshold, indicating that nuclear interactions may affect the cell dynamics and structure in these tissue layers.

While both cell and nuclear anisotropies as well as shape factors were large at early developmental stages of the retina, when the tissue features a neuroepithelial architecture (Fig. 4d; Methods; Extended Fig. 4a,b), they decreased progressively over time as cells detached from tissue boundaries and continued doing so after that (Fig. 4g). As the nuclear volume fraction progressively reaches very high levels after cell detachment from tissue boundaries (55-60 hpf; Fig. 4a,d), with nearly the whole cell occupied by the nucleus, nuclei started to display shape deformations associated with the strong nuclear jamming conditions (Fig. 4b-b’). Eventually, as development progressed, both cell bodies and nuclei displayed more compact shapes, with nuclei becoming nearly round in both the INL and RGL (Fig. 4g; *𝛼*_*N*_ ≃ 1.2 at 120 hpf).

Since our theoretical predictions indicated that jamming of round nuclei can enhance the order of cellular packings (Fig. 2g-k), we analyzed cellular order in the INL. From the network spanned by connections between the centers of nuclei’s nearest neighbors (Fig. 4h), we obtained both the lengths between a nucleus and its nearest neighbors (bond lengths; Methods) as well as the angles between nearest neighbors (bond angles; Methods) at both 55 hpf and 120 hpf (Fig. 4h). We observed that the variations in both bond lengths (Fig. 4i) and angles (Fig. 4j) showed significant reductions from 55 hpf to 120 hpf. These results indicate that the cellular packing structure of the INL becomes more ordered as the jammed nuclei become rounder, with cells forming a nearly crystalline arrangement at 120 hpf, as predicted by our simulations. This is consistent with the formation of the precise and almost crystalline ordering of the neighboring retinal cell-types known as retinal mosaics, which is proposed to ensure the even distribution of retinal cells and thereby tissue functionality^4,50,51^.

## Nuclear jamming in brain tissues

Beyond eye development, we examined other neural tissues to understand if a similar behavior was observed. In particular, we compared the characteristics of cell and nuclear sizes and shapes in the midbrain (MB) at early (18 hpf) and later (92 hpf) stages of brain development in zebrafish^52^. At 18 hpf, the MB architecture is that of a pseudostratified neuroepithelium (Fig. 5a-a’’’’). After cells detach from the tissue boundaries, the dorsal part of the MB later gives rise to the optic tectum (OT), a tissue which coordinates eye movement and visual processing in zebrafish and other amphibians^53^ (Fig. 5b-b’’’’). Our analysis showed that both cell and nuclear sizes as well as their aspect ratios strongly decreased from their values in the MB (18 hpf) to those measured in the OT (92 hpf), with cell size and aspect ratio decreasing more than nuclei’s (Fig. 5c,d), as was the case in the retina. While the measured nuclear volume fraction is below the nuclear jamming threshold in the MB (Fig. 5e), it is larger than the threshold in the OT, indicating that the tissue underwent a nuclear jamming transition during or after cells detached from tissue boundaries. In addition, the shape factor of both cells and nuclei strongly decreased from the MB to the OT (Fig. 5f), as cells become more compact (Fig. 5b’’’’) and nuclei nearly round (Fig. 5b’’’). Overall, these data show that nuclear jamming also occurs in these brain tissues, with very similar characteristics as those reported above for retina development, including the more ordered cell packings (Fig. 5b’’’) associated with the jamming of round nuclei.

**Figure 5.**
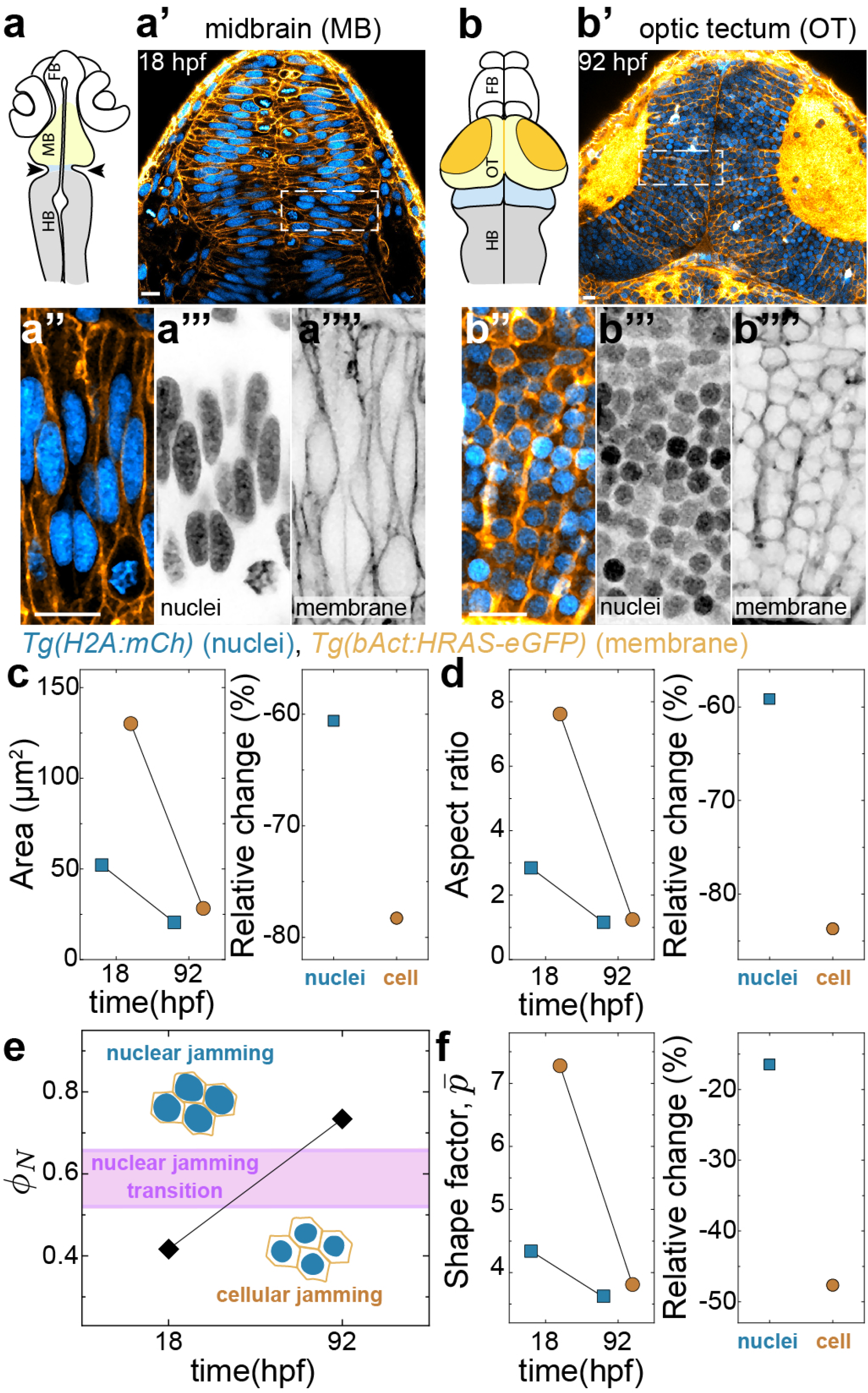
Nuclear jamming in brain development. **a**, Schematic representation of a 18 hpf zebrafish, depicting the midbrain (MB, light yellow), hindbrain (HB, gray) and forebrain (FB, white), with arrows indicating the MB-HB boundary (blue). **a’**, Representative confocal section of a 18 hpf zebrafish midbrain (nuclear label, *Tg(H2A:mCh)*, cyan; membrane label, *Tg(bAct:HRAS-eGFP)*, orange). Scale bars, 10 μm. **a’’-a’’’’**, Higher magnification insets of the outlined region in a’. **b**, Schematic representation of a 92 hpf zebrafish. MB developed into the optic tectum (OT, light yellow). **b’**, Representative confocal section of a 92 hpf zebrafish OT (same markers as in a’). **b’’-b’’’’**, Higher magnification insets of the outlined region in b’. Scale bars, 10 μm. **c-d**, Areas (c) and aspect ratios (d) of cells (orange) and nuclei (blue) in the MB (18 hpf) and OT (92 hpf). Their relative change between 18 to 92 hpf is plotted on the right panels. **e**, Nuclear volume fraction in the MB (18 hpf) and in the OT (92 hpf). Horizontal violet band shows the nuclear jamming volume fraction predicted theoretically (Fig. 2). **f**, Shape factor of cells (orange) and nuclei (blue) in the MB (18 hpf) and OT (92 hpf). Their relative change between 18 to 92 hpf is plotted on the right panels. MB (N=6, n=312), OT (N=6, n=419).

## Phase diagram of nuclear jamming

To understand the different phases of nuclear jamming, we computed the phase diagram as a function of the two relevant parameters in the system, namely the nuclear volume fraction *𝜙*_*N*_ and nuclear anisotropy *𝛼*_*N*_. For small values of nuclear volume fraction, the system displays cellular jamming but not nuclear jamming (Fig. 6). As nuclear volume fraction increases over an anisotropy-dependent critical volume fraction 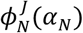, the system undergoes either a transition to jammed isotropic nuclei for small (or no) nuclear anisotropy or a transition to jammed anisotropic nuclei without strongly affecting cell shapes. Large nuclear anisotropy and volume fraction levels lead to strong jamming of anisotropic nuclei that substantially deform cell shapes, causing local nematic ordering in the system.

**Figure 6.**
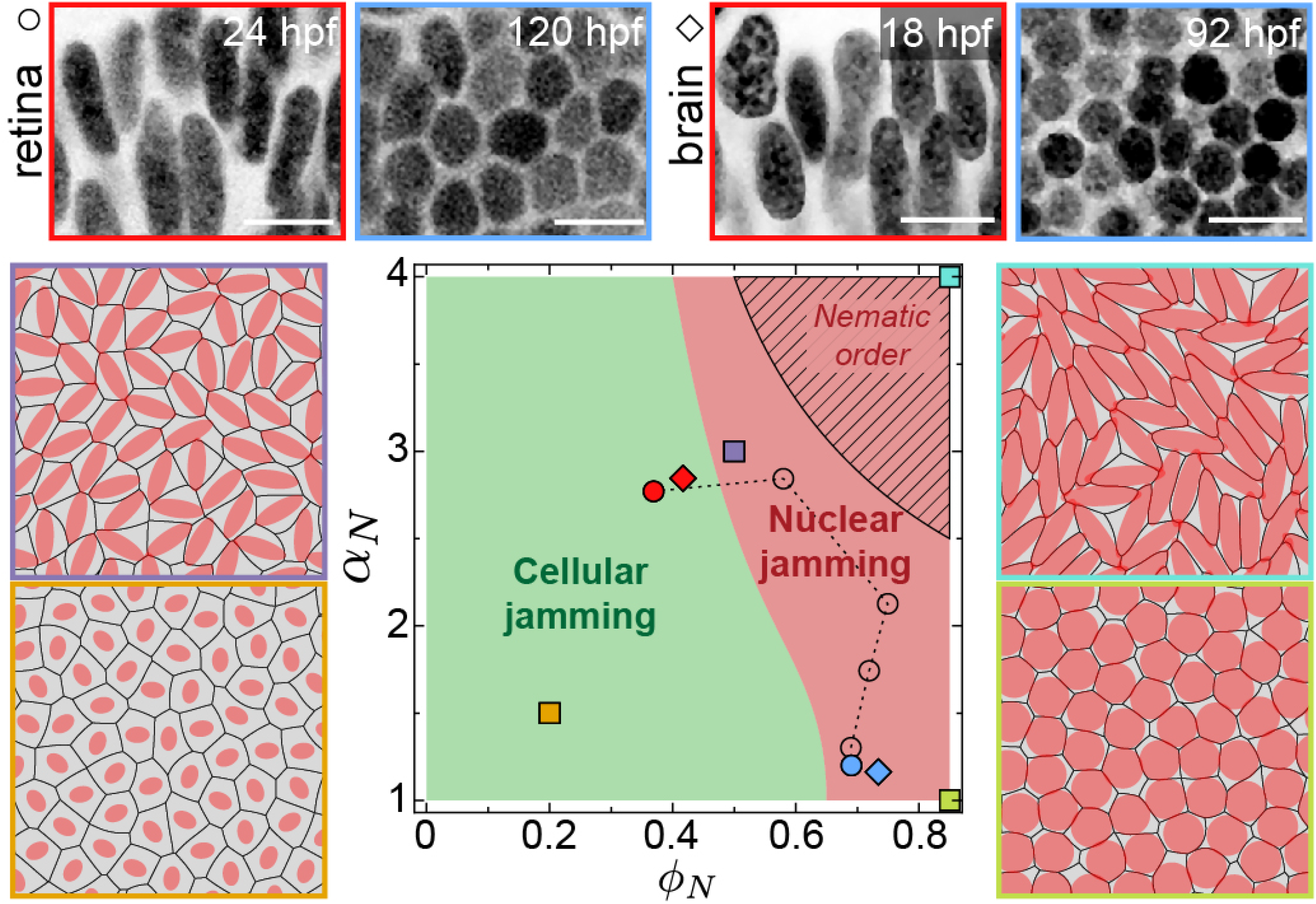
Phase space for nuclear jamming. Predicted phase space for varying values of the nuclear volume fraction 𝜙_*N*_and the nuclear anisotropy 𝛼_*N*_. Representative simulation snapshots for limiting phases are shown (orange: cellular jamming, but no nuclear jamming; green: nuclear jammed state for round nuclei; violet: onset of anisotropic nuclear jamming; cyan: nuclear jammed states with local nematic order). Dashed region indicates existence of local nematic order. The developmental trajectory (dashed line) of the INL tissue in the retina (circles) in the phase space is shown, from 24 hpf (red circle) to 120 hpf (blue circle). The location of the phase space of the midbrain at 18 hpf (red diamond) and after developing into the optic tectum (92 hpf; blue diamond) are also shown. Representative confocal sections of nuclei in the retina at 24 hpf (red) and in the INL at 120 hpf (blue), as well as in the midbrain (18 hpf; red) and optic tectum (92 hpf; blue) are shown (top).

Plotting the measured developmental trajectory for the retina (INL) in the phase space, we observe that cells in the retina first undergo nuclear jamming with highly anisotropic nuclei and subsequently decrease their nuclear aspect ratio maintaining the nuclear jammed state. This progressive reduction of nuclear anisotropy in a jammed nuclear state causes the observed ordering of cellular packings, eventually generating a nearly crystalline structure. While we lack an exhaustive developmental trajectory for brain tissues, the initial MB state (18 hpf) and final OT state (92 hpf) are essentially located in the same region of the phase space as the initial and final stages of retina development, suggesting that different neural tissues may follow similar developmental trajectories as they undergo nuclear jamming.

## Discussion

Understanding the role of subcellular structures in the control of the emergent tissue physical state and architecture is essential to connect relevant biological processes occurring at different scales during embryonic development. Our computational and experimental results show that nuclei play an important role in the emergent physical state of the tissue during organ formation in vertebrates.

We introduced nuclei in Active Foam description, enabling studies of the interplay between nuclear properties and cell and tissue structure and dynamics. In the absence of nuclei, previous models predicted the existence of rigidity transitions in multicellular systems^7,27,32,35^, such as embryonic tissues. Our computational results show that large values of the nuclear volume fraction cause nuclear jamming, which physically affects tissue architecture by strongly reducing cellular movements and T1 transitions as well as increasing tissue rigidity and cellular order. In this sense, nuclear jamming is a rigidity transition occurring in the presence of different rigidity transition, namely cellular jamming. By controlling the nuclear volume fraction, cells can change the emergent properties of the tissue from being controlled by the mechanics of cell-cell contacts to the mechanics of the nuclei, as well as change the structure of cellular packings, linking directly the physical properties of the nucleus to the physical state of the tissue.

Our experimental data indicates that nuclear jamming occurs during the development of both eye and brain tissues in zebrafish. We observed a progressive increase in the nuclear volume fraction over time, eventually crossing the nuclear jamming threshold. For large values of the nuclear volume fraction, we observed that the structure of cellular packings becomes more ordered, as predicted theoretically, resembling the nearly crystalline cellular packings in the adult functional eye. Nuclear jamming thus provides a different cell ordering mechanism than those reported for epithelial tissues^3,4^. In a nuclear jammed state, high nucleus stiffness translates into a stiffer tissue, which could also have a functional role by providing further mechanical integrity to the tissue.

These results highlight the importance of nuclear volume fraction in the control of the emergent tissue physical state and architecture, and provide a direct connection between the supracellular state and a subcellular organelle. Future studies will tell if nuclear jamming occurs in other organs and species, as well as establish its functional roles.

## Data Availability Statement

The data that supports these findings are available upon request.

## Acknowledgements

We thank Claudia Froeb, Georgina Stooke-Vaughan and Arthur Boutillon for technical support. We also thank the UCSB Animal Research Center and both the animal facility and the imaging facility at MPI-CBG for their support. This work was supported by the Eunice Kennedy Shriver National Institute of Child Health and Human Development of the National Institutes of Health (R01HD095797), and the Deutsche Forschungsgemeinschaft (DFG, German Research Foundation) under Germany’s Excellence Strategy – EXC 2068 – 390729961– Cluster of Excellence Physics of Life of TU Dresden. We acknowledge support from the Center for Scientific Computing from the CNSI, MRL: an NSF MRSEC (DMR-1720256) and NSF CNS-1725797.

## Author Contributions

SK, RA and OC designed research; RA performed all experiments; SK and RA analyzed the data; SK performed all simulations; SK, RA and OC wrote the paper; OC supervised the project.

## Competing Financial Interests Statement

The authors declare that they have no competing financial interests.

## FIGURE LEGENDS

**Extended Figure 1.**
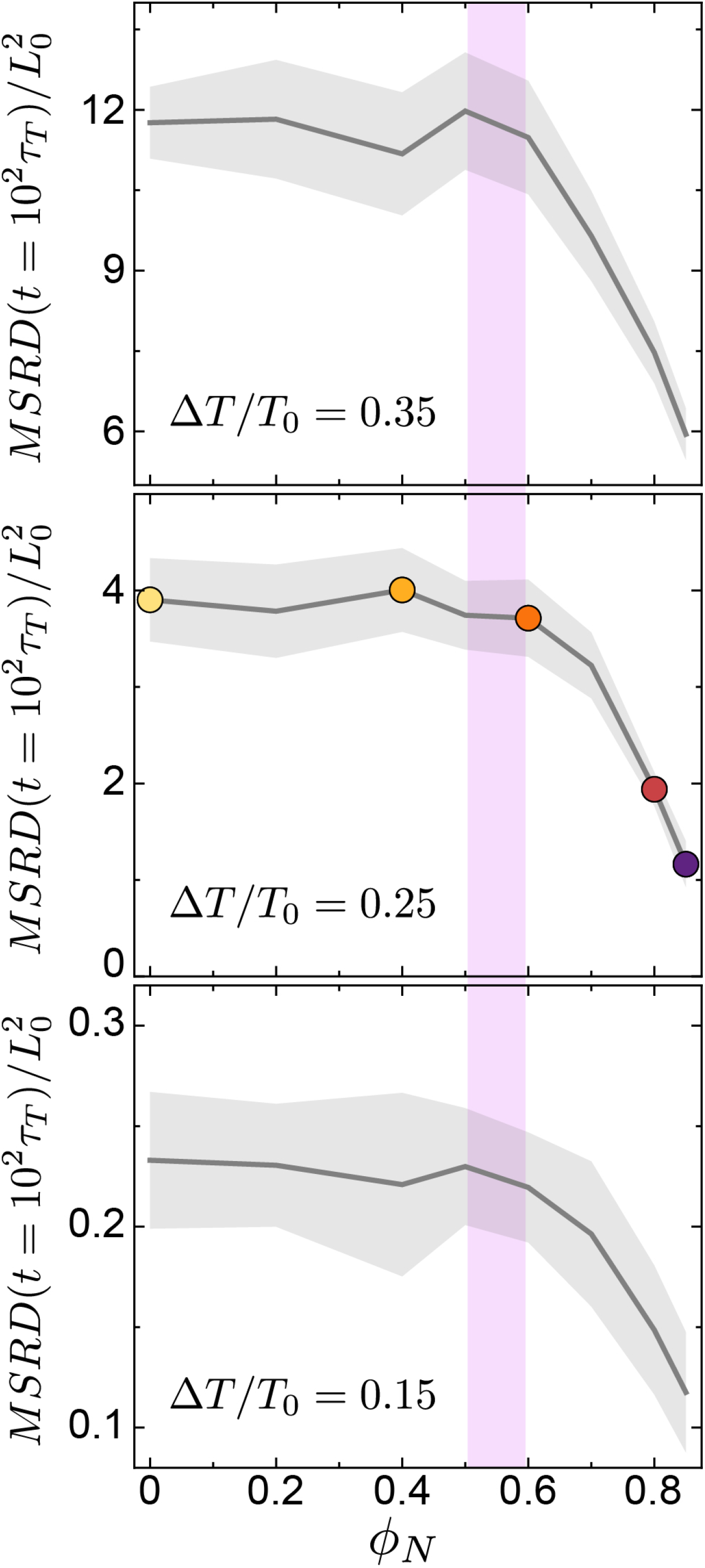
Effects of the magnitude of tension fluctuations on the nuclear jamming transition. Dependence of long timescale MSRD values in terms of nuclear volume fraction for different magnitudes of tension fluctuations Δ*T*/*T*_0_, showing that the dynamic slowdown in cell movements starts to occur at the same nuclear volume fraction (violet band) regardless of tension fluctuation strength.

**Extended Figure 2.**
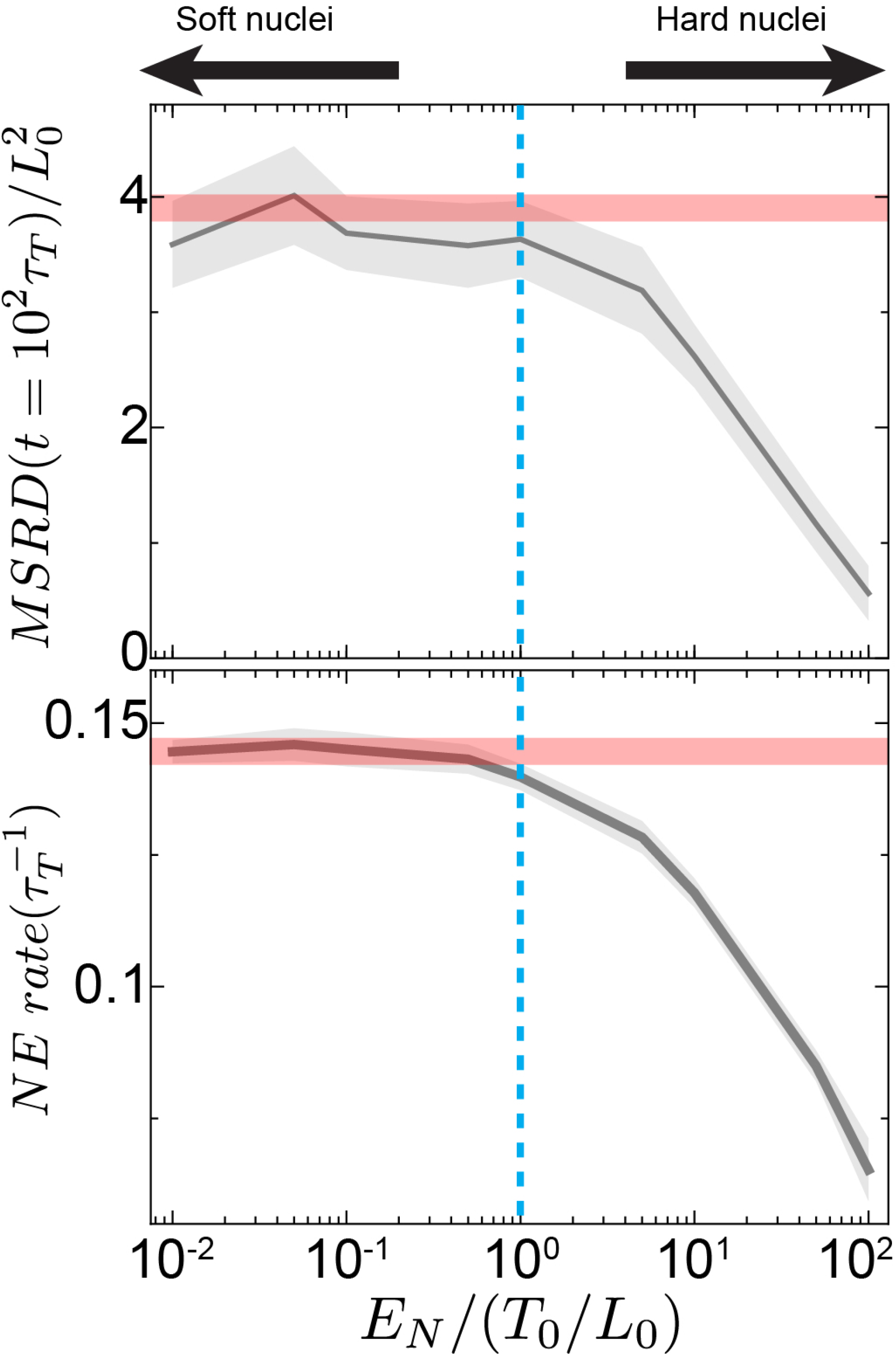
Effects of nuclear stiffness on the nuclear jamming transition. Dependence of the long timescale MSRD values (top) and neighbor exchange rate (bottom) for large nuclear volume fraction (*𝜙*_*N*_ = 0.85) on the ratio of nuclear stiffness *E_N_*and the characteristic stress scale of cell deformations *T*_0_/*L*_0_. Large (small) values of *E_N_*/(*T*_0_/*L*_0_) indicate hard (soft) nuclei compared to the cell. Red lines are the values of long time MSRD and neighbor exchange rate for zero nuclear volume fraction (𝜙_*N*_ = 0), and the vertical cyan dashed line indicates the point at which nuclear stiffness and the stress scale of cell deformations are equal, namely *E_N_* = *T*_0_/*L*_0_.

**Extended Figure 3.**
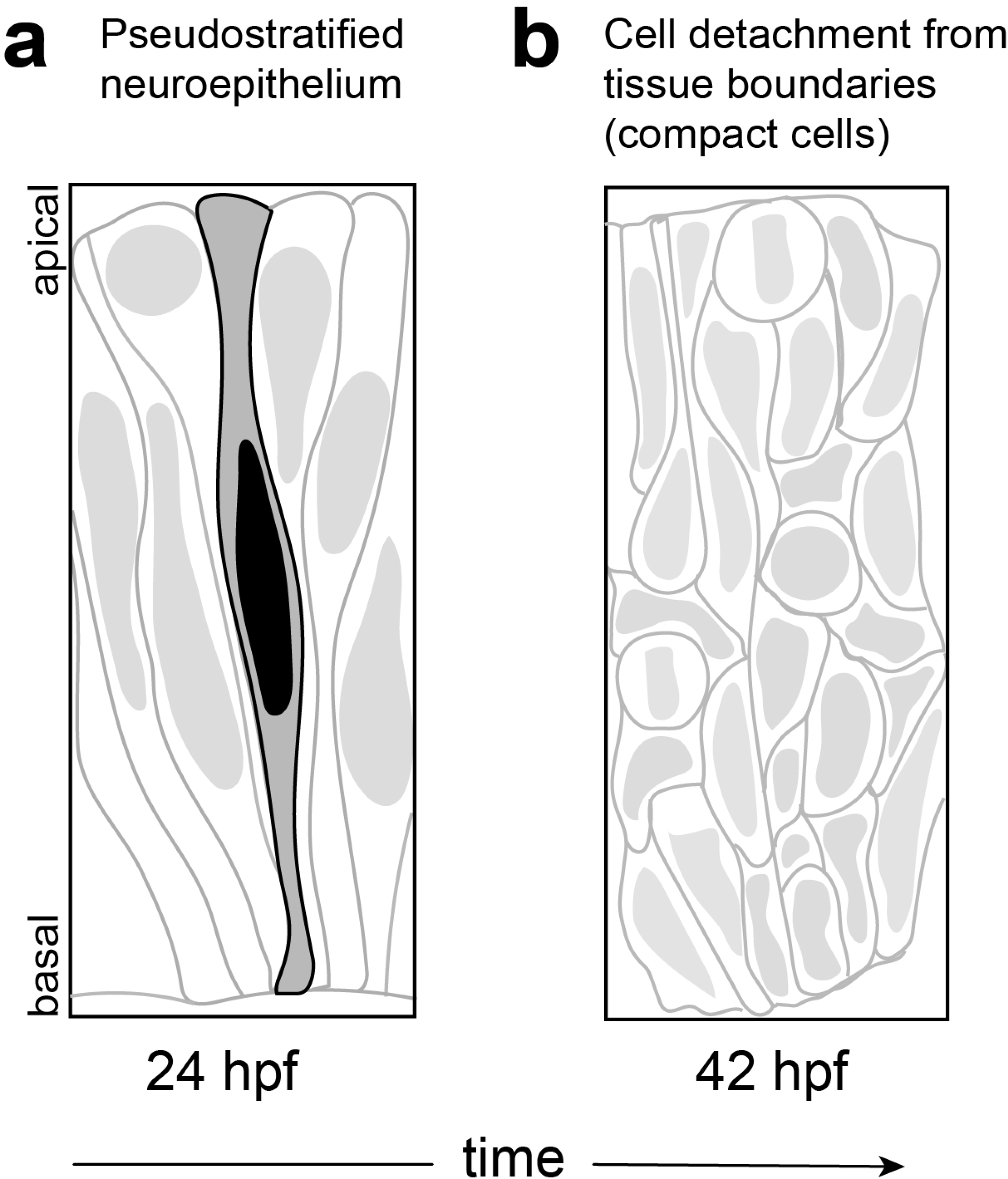
Schematic representation of the zebrafish retina before and after cell detachment from apical and basal tissue boundaries. **a**, The zebrafish is a single-layered neuroepithelium with elongated cells connected to both the apical and basal tissue surfaces at 24 hpf. **b**, At approximately 40-42 hpf, majority of retinal cells detach from the tissue (apical and basal) boundaries and form a tissue with compact cells. This single-layered tissue eventually evolves into the different layers described in the main text (Fig. 4).

**Extended Figure 4.**
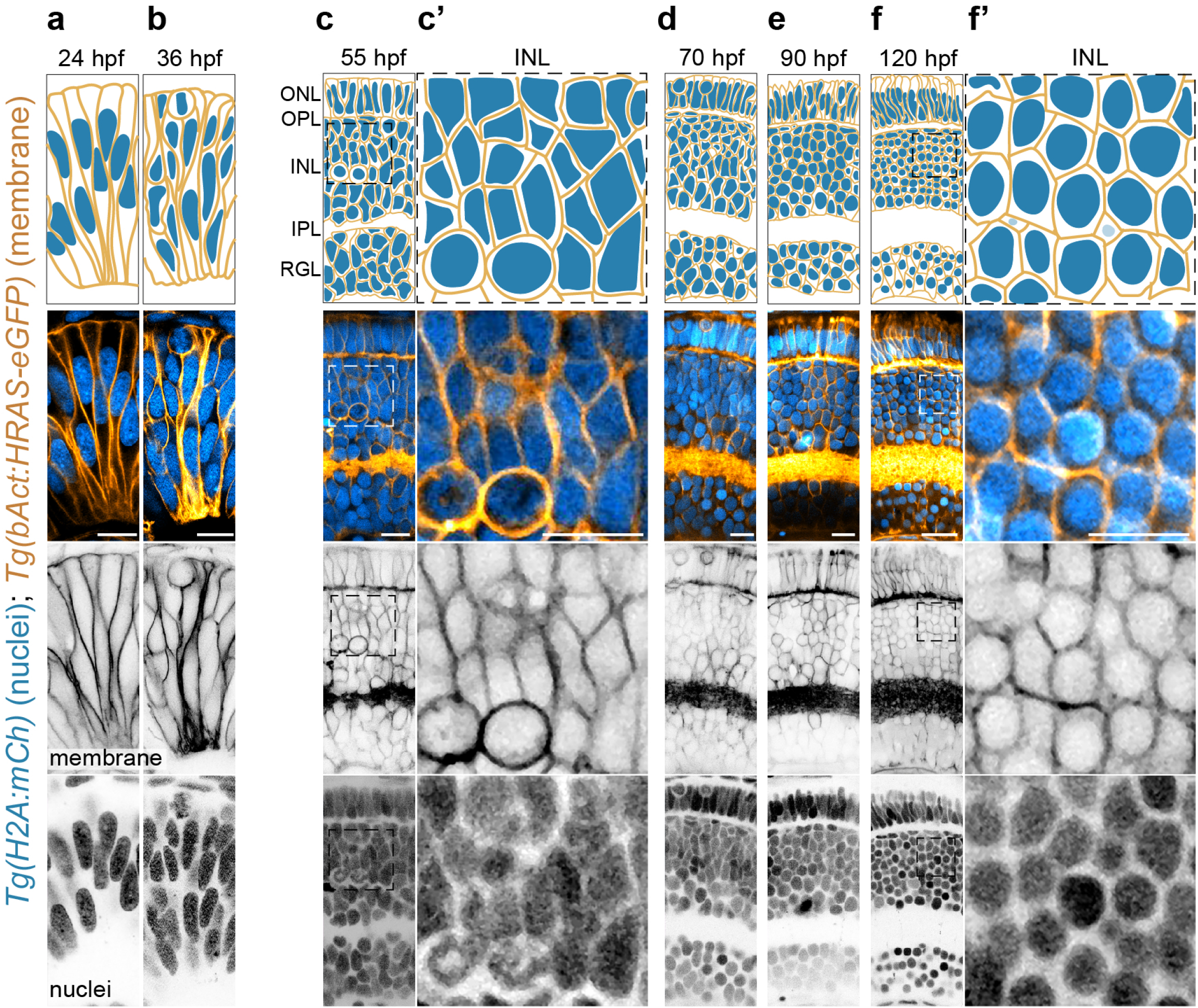
Tissue architecture during retina development: evolution of cell and nuclear shapes and sizes. a-f’, Top: schematic representation of cellular and nuclear architecture during different retinal stages (a, 24; b, 36; c, 55; d, 70; e, 90; f, 120 hpf). Bottom: Representative confocal sections of zebrafish retinae expressing the membrane marker (*Tg(bAct:HRAS-eGFP)*; orange) and nuclei marker (*Tg(H2A:mCh)*; cyan). Scale bar, 20 μm. **c’**, Higher magnification inset of the outlined region in **c. f’**, Higher magnification inset of the outlined region in **f**. Scale bars, 10 μm.

## ONLINE METHODS

### Zebrafish husbandry, fish lines, and labeling

Zebrafish (*Danio rerio*) were maintained as previously described^54^. Experiments were performed following all ethical regulations and according to protocols approved by the European Union (EU) directive 2010/63/EU as well as the German Animal Welfare act. For ubiquitous labeling of cell membranes we used *Tg(bAct:HRAS-EGFP)* embryos (Cooper et al. 2005). For ubiquitous nuclear labelling we used *Tg(H2A:mCh)* (Knopf et al. 2011).

### Imaging

Embryos were manually dechorionated and anesthetized in 0.02% tricane methanesulfonate (MS-222; Sigma-Aldrich) in E3 medium for approximately 20 minutes. The medium was supplemented with 0.2 mM 1-phenyl-2-thiourea (PTU) (Sigma-Aldrich) from 8±1 hpf onwards to prevent pigmentation. Immobilized embryos were then placed in 1% low-melting-point agarose (in E3) in glass-bottom dishes (MatTek Corporation). Samples were mounted for lateral or dorsal imaging for retina and brain imaging, respectively. The dishes were filled with E3 medium containing MS-222 (0.02%) and 1-phenyl-2-thiourea (PTU, 0.003%, Sigma-Aldrich) and imaged at room temperature.

Samples from 18-70 hpf were imaged on a Laser Scanning Confocal Microscope (Zeiss, LSM780) using the 40x/1.2 C-Apochromat water immersion objective (Zeiss). Z-stacks were recorded with 0.44 μm confocal sectioning. For later developmental stages with high tissue thickness (90-120 hpf), we used a Multiphoton Laser Scanning Confocal Microscope (2-photon inverted microscope; Leica, SP8) with a 40x/1.1 HC PL IRAPO, water immersion objective (Leica). Z-stacks were recorded with 0.5 μm optical sectioning.

### 2D Analysis of the cells and nuclei sizes and shapes

To reduce noise in original images, we used ImageJ’s 3D median filter of 1 pixel radius. We first detected the middle of each nucleus in 3D by finding their maximal cross-sectional area in the 3D stack. Then, we used the polygon selections tool in ImageJ to trace the outline of each cell and nucleus at z-plane corresponding to the 3D middle of each nucleus. Using *Fit Ellipse* tool in ImageJ, we next extracted the following shape descriptors: Area, major axis (primary axis of the best fitting ellipse), minor axis (secondary axis of the best fitting ellipse), Aspect Ratio (AR) (major axis/minor axis) and circularity, where the circularity is used to compute the shape factor in the analysis.

### 2D Analysis of the nuclei ordering

We manually identify nuclei of retinal cells within the INL at 10 retinae at 55 hpf and 10 retinae at 120 hpf. For each sample, nearest neighbor relations were computed using Voronoi tessellation based on nuclei positions. The network configuration was constructed by connecting all pairs of nearest neighbors. The bond length, *s*, and the bond angle, *𝜃*, are defined in the same way as in the analysis of tissue structure in simulations (see above; Fig. 2j,k). The deviation of tissue structure from the crystalline hexagonal structure was quantified by the standard deviation of the normalized bond length 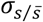 and bond angle 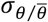.

### Theoretical description

We extended the previously published Active Foam computational framework for tissue dynamics to include nuclei. Tangential and normal forces at cell boundaries (accounting for cortical tension and adhesion, as well as osmotic pressure), nucleus-cell boundary interaction forces and a nucleus restoring force towards the geometric cell center were all implemented (Fig. 2c; Supplementary Information). The effective interaction potential between nuclei and cell boundaries was approximated as a repulsive harmonic potential, with the resulting repulsive force being along the normal direction of the cell boundary at every point of interaction. The total repulsive force acting on each nucleus is the integral of the distributed interaction forces along the cell boundary. The restoring force for the nucleus to cell center is modelled as a linear force acting on the nucleus center and pulling it towards the geometric cell center. See Supplementary Information for details.

### Simulations

All simulations included *N* = 256 cells in a periodic square box. An initial configuration consisting of a random Poisson Voronoi tessellation was first generated in a periodic square box and the system was allowed to relax for 10 *τ_T_* before sampling data points. Confluency (no spaces between cells) was enforced for all simulations. The governing equations are non-dimensionalized with characteristic scales (Supplementary Information), and they are numerically integrated using the Euler method and the Euler-Maruyama method with a time step, Δ*t* = 0.01*τ_R_*, where *τ_R_* is the vertex relaxation time scale that is the smallest characteristic timescale in the system (Supplementary Information).

### Analysis of simulated cell trajectories and MSRD

Cell trajectories were computed by tracking the center positions of all cells for each time point. To quantify cell movements, we computed the mean-squared relative displacement (MSRD), as previously done^6^. For each cell trajectory *r_i_*(*t*), the nearest neighbor at an initial time point was first identified and a relative distance vector between cell i and its nearest neighbor cell j, *r_ij_*(*t*) = *r_i_*(*t*) – *r_j_*(*t*), was computed accordingly. We computed the mean-squared relative displacement of a given cell i, M*SRD_i_*(*t*) = (*r_ij_*(*t*) – *r_ij_*(0))^2^, and MSRD values were averaged over all cells as well as all initial time points. MSRD values were normalized by the characteristic length scale of cell size, *L*_0_.

### Analysis of tissue structure in simulations

To investigate structural changes in simulated tissues, we used four different structure measures: (1) shape factor, (2) defect density, (3) bond length variation, and (4) bond angle variation. The shape factor 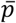 was defined as mean of the ratio of cell perimeter *P_i_* to square root of cell area *A_i_*, namely

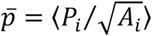

This measure quantifies the degree of anisotropy in cell shape and was previously used to identify density-independent rigidity transition^24,32^. The defect density 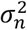 was defined as the variance of the topology distribution and it quantifies number density of cells that have a number of neighbors different from six. With *n*_/_ being number of neighbors for cell I, the defect density can be computed as

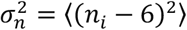

A network structure of a given tissue configuration (cellular packings) was constructed by connecting bonds between all cells with their neighbors. The bond length *s* is the length of the line connecting a cell and a nearest neighbor (Fig. 2j, inset) and the bond angle *𝜃* is the angle between two adjacent nearest neighbors of a given cell (Fig. 2k, inset). The standard deviation of normalized bond length 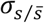 and that of normalized bond angle 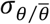 were computed accordingly to quantify the degree of variation in these measures.

In two dimensional confluent systems, the ground state of the monodisperse cell packing exhibits crystalline hexagonal order. This situation corresponds to a minimum shape factor for the confluence condition, zero defect density, and zero variation in the bond length and the bond angle, respectively. Hence, any deviation from the ground state structural measures quantifies the degree of structural disorder in the system.

To quantify local nuclei alignment, we defined the local nematic order parameter as follow. For a given pair of neighboring cell i and cell j with nuclei orientation *𝜃_i_* and *𝜃_j_*, their nuclei alignment is computed as cos 2(*𝜃_i_* – *𝜃_j_*), where the value of 1 and -1 correspond to the same orientation and the perpendicular orientation of neighboring nuclei, respectively. The local nematic order parameter 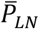 is equal to the mean of the alignment measure over all neighboring pairs of cells, namely

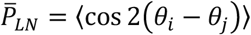

The local nematic parameter value is positive when neighboring nuclei align in the same direction while the random orientation of nuclei corresponds to zero.

To compute the mean number of contacts between nuclei and cell boundaries, 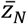, all cell boundaries of a given cell that exert repulsive force to its nucleus were counted as contacting cell boundaries. Taking the average of number of contacting cell boundaries over all cells, we can quantify the degree of interaction between nuclei and cell boundaries.

## Notes

### Competing Interest Statement

The authors have declared no competing interest.

