## Supplementary Information for "A nuclear jamming transition in vertebrate organogenesis"

### Supplementary Information to
### *A nuclear jamming transition in vertebrate organogenesis*

Sangwoo Kim, Rana Amini, and Otger Campàs

#### Captions of Supplementary Movies

##### Supplementary movie 1

Simulations of the system dynamics for isotropic nuclei with varying nuclear volume fraction ($\phi_N = 0.2, 0.5$, and 0.85 from left to right). Trajectories of sixteen cells are shown in different colors.

##### Supplementary movie 2

Simulations of the system dynamics for varying nuclear aspect ratio at large volume fraction ($\alpha_N = 1, 2, 3$, and 4 from left to right and $\phi_N = 0.85$). Trajectories of sixteen cells are shown in different colors.

### Theoretical Description of Nuclear Jamming: Governing Equations

Building up on the existing Active Foam description of tissue dynamics (Ref. 7 in the main text), we describe cells via a vertex model that includes tension dynamics at the cell-cell contacts, and introduce nuclei and their dynamics to the Active Foam description. Tissue dynamics and structure are therefore described by movements of cell vertices and nuclei. The degrees of freedom in the system are the triple vertices (physical vertices), the intermediate vertices (non-physical vertices), as well as nuclei positions and orientations (Fig. 2a of main text). The dynamics of vertices and nuclei result from force balance in an overdamped environment, namely

$$\eta_R \frac{d\boldsymbol{R}_\alpha}{dt} = \sum_{i,j \in F(\alpha)} \left( \boldsymbol{T}_{ij} \Theta\left(T_{ij}\right) + \boldsymbol{N}_{ij} \right) + \sum_{i \in F(\alpha)} , \boldsymbol{F}_{\alpha i} \tag{1}$$

$$\eta_N \frac{d\boldsymbol{R}_{N,i}}{dt} = - \sum_{\alpha \in V(i)} \boldsymbol{F}_{\alpha i} + \boldsymbol{G}_i \tag{2}$$

$$\eta_\theta \frac{d\theta_{N,i}}{dt} = - \sum_{\alpha \in V(i)} \boldsymbol{R}_{\alpha i} \times \boldsymbol{F}_{\alpha i} = M_i \tag{3}$$

Here, $\boldsymbol{R}_\alpha$, $\boldsymbol{R}_{N,i}$, and $\theta_{N,i}$ are the position of vertex $\alpha$, the position of nucleus of cell $i$, and the orientation of nucleus of cell $i$, respectively, and $t$ is time. $\eta_R$, $\eta_N$, and $\eta_\theta$ are friction coefficients of vertex position, nucleus position, and nucleus orientation, respectively. As in the previous Active Foam description (Ref. 7 in the main text), $\boldsymbol{T}_{ij}$ is the effective tension at the cell junction between cell $i$ and $j$, $\boldsymbol{N}_{ij}$ is the normal force acting on vertex $\alpha$ (Fig. S1a), $\boldsymbol{F}_{\alpha i}$ is the repulsive force between nucleus of cell $i$ and vertex $\alpha$ (Fig. S1b), and $\boldsymbol{G}_i$ is a restoring force acting on the nucleus, directed to the geometric center of a given cell (Fig. S1c). $\boldsymbol{R}_{\alpha i}$ is the distance between the nucleus of cell $i$ and vertex $\alpha$. $\Theta(\cdot)$ is the Heaviside step function that prevents unrealistic negative tensions. $F(\alpha)$ represents all cells sharing vertex $\alpha$ and $V(i)$ represents all vertices sharing cell $i$.

As in the Active Foam description, the dynamics of tensions $T_{ij}$ are those of an Ornstein-Uhlenbeck process, with the effective tension fluctuating around a fixed tension $T^0_{ij}$ and a persistent time $\tau_T$ for tension changes, namely

$$\tau_T \frac{dT_{ij}}{dt} = -(T_{ij} - T^0_{ij}) + \Delta T \sqrt{2\tau_T}\xi\,, \tag{4}$$

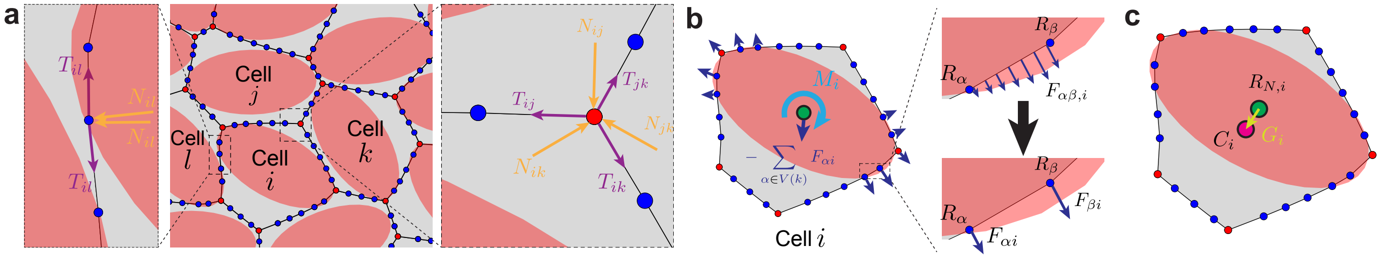


**Figure S1: Effective forces acting on vertices and nucleus.** **a** Schematics of tangential and normal forces acting on an intermediate vertex (left) and triple vertex (right). **b** Schematics of the repulsive interaction between nuclei and cell junctions. **c** Schematics of the nucleus restoring force.

where $\Delta T$ is the magnitude of tension fluctuations and $\xi$ is Gaussian white noise. Since all simulations in this work focus on the confluent regime with identical cell types, the fixed tension value, $T_{ij}^0$, is set to be same for all cells, $T_0$. The magnitude of the normal forces, $N_{ij}$, arises from the osmotic pressure difference between adjacent cells and is given by $N_{ij} = \left(\Delta P_i - \Delta P_j\right) L_{ij}/2$, where $L_{ij}$ and $\Delta P_i (\Delta P_j)$ are the contour length of junction between cell $i$ and $j$ and the osmotic pressure difference across cell $i(j)$, respectively. The osmotic pressure difference is shown to be inversely proportional to cell size so it can be expressed as $\Delta P_i = K/A_i - P_0$, where $K$ is the cell compressibility and $P_0$ is the osmotic pressure outside cells, which is set to be constant.

The interaction between cell boundaries and nuclei is modeled as a repulsive harmonic interaction. A repulsive force and torque, $F_{\alpha\beta,i}$ and $M_{\alpha\beta,i}$ respectively, act on the cell boundary $\alpha\beta$ between vertex $\alpha$ and $\beta$ by the nucleus of cell $i$, and are determined by the integral of the distributed forces along the cell junction, namely

$$F_{\alpha\beta,i} = \int_0^{L_{\alpha\beta}} K_n \left(R_{0,i}(x) - R_i(x)\right) \Theta\left(R_{0,i}(x) - R_i(x)\right) dx, \tag{5}$$

$$M_{\alpha\beta,i} = \int_0^{L_{\alpha\beta}} \left(x - \frac{L_{\alpha\beta}}{2}\right) \times K_n \left(R_{0,i}(x) - R_i(x)\right) \Theta\left(R_{0,i}(x) - R_i(x)\right) dx, \tag{6}$$

where $K_n \equiv E_N/L_0$ (with $E_N$ being the nuclear stiffness), $R_{0,i}(x)$ is the length of the nucleus axis in the direction connecting the nucleus center and an intermediate point $x$ along cell junction $\alpha\beta$, and $R_i(x)$ is the distance between the nucleus center and the intermediate point $x$. The effective force and torque can be replaced by point forces acting on two end vertices, vertex $\alpha$ and vertex $\beta$, namely, $F_{\alpha i}$ and $F_{\beta i}$. The resulting force acting on the nucleus $i$ from all cell junctions $\alpha\beta$ is simply sum over all reaction forces. The resulting moment can computed by summing over cross products of a vector connecting nucleus center to an individual vertex and the reaction force for all vertices (Fig. S1b).

Finally, the restoring force $\boldsymbol{G}_i$ on the nucleus of cell $i$ is assumed to be proportional to the distance of the nucleus center from the cell center and in the direction connecting the nucleus center to the cell center (Fig. S1c), and reads

$$\boldsymbol{G}_i = K_c \left(\boldsymbol{C}_i - \boldsymbol{R}_{N,i}\right), \tag{7}$$

where $K_c$ is a restoring force modulus and $\boldsymbol{C}_i$ and $\boldsymbol{R}_{N,i}$ are the cell and nucleus center positions of a given cell $i$, respectively.

#### Dimensionless Parameters

The above equations can be made dimensionless normalizing by the characteristic scales, namely the effective tension scale $T_0$, the cell size length scale $L_0 = \sqrt{A_0} = \sqrt{K/P_0}$, and the vertex relaxation time scale $\tau_R = \eta_R L_0/T_0$. After normalizing all variables, we obtain all dimensionless parameters, which are summarized in the Table below.

| | |
|---|---|
| $\phi_N = A_N/A_0$ | Nuclear volume fraction: ratio of nucleus size to cell size |
| $\alpha_N = b/a$ | Nuclear aspect ratio: ratio of the major semiaxis $b$ to the minor semiaxis $a$ of the nucleus |
| $\Delta T/T_0$ | Magnitude of tension fluctuations |
| $\tau_T/\tau_R$ | Ratio of tension relaxation timescale to vertex relaxation timescale |
| $\tau_N/\tau_R$ | Ratio of nucleus translation relaxation timescale to vertex relaxation timescale |
| $\tau_\theta/\tau_R$ | Ratio of nucleus rotational relaxation timescale to vertex relaxation timescale |
| $P_0L_0/T_0$ | Ratio of normal force scale to stress scale of cell deformations |
| $E_N/(T_0/L_0)$ | Ratio of nuclear stiffness to stress scale of cell deformations |
| $K_cL_0/T_0$ | Ratio of nucleus restoring force scale to junctional tension scale |

We assume that the timescales relevant to nucleus relaxation are comparable to that of vertex relaxation so that $\tau_N/\tau_R$ and $\tau_\theta/\tau_R$ are set to be 1 while $\tau_T/\tau_R$ is set to be 10 as the tension relaxation time scale is experimentally reported to be longer than the vertex relaxation time scale. $P_0L_0/T_0$ is set to be 10 as the osmotic pressures are expected to be larger than cortical tension. Nucleus is known to be much stiffer compared to the rest of cytoplasm as well as cell membrane so $K_nL_0^2/T_0$ is set to be 50. $K_c$ is set to be 1 to ensure that the nucleus center is always close to the geometric cell center.
